# Profiling of Human Gut Virome with Oxford Nanopore Technology

**DOI:** 10.1101/2020.02.03.933077

**Authors:** Jiabao Cao, Yuqing Zhang, Min Dai, Jiayue Xu, Liang Chen, Faming Zhang, Na Zhao, Jun Wang

**Author notes:** Jiabao Cao, and Yuqing Zhang contributed equally to this manuscript. Correspondence to: Professor Jun Wang, CAS Key Laboratory of Pathogenic Microbiology and Immunology, Institute of Microbiology, Chinese Academy of Sciences, No. 1-3 Beichenxi Road, Chaoyang District, Beijing, China; Dr. Na Zhao, CAS Key Laboratory of Pathogenic Microbiology and Immunology, Institute of Microbiology, Chinese Academy of Sciences, No. 1-3 Beichenxi Road, Chaoyang District, Beijing, China;.

## Abstract

Human gut virome play critical roles in maintaining gut microbial composition and functionality, as well as host physiology and immunology. Yet, there are insufficient amount of studies on this topic mainly due to methodological limitations, including enrichment of viruses (phages and host viruses) as well as short read-length from current sequencing technology. Here we developed a full working protocol for analyzing human gut virome using physical enrichment, reverse transcription and random amplification, and eventually the state-of-art single-molecule real-time sequencing (SMRT) platform of Oxford Nanopore Technology (ONT). We demonstrate that sequencing viral DNA directly, or viral DNA/cDNA after amplification using ONT achieves much longer reads and provides more information regarding virome diversity, many of the virome sequences do not have match in current databases. Moreover, direct DNA sequencing of virome provides first insights into the epigenetic modifications on phages, where signals of methylations can be directly detected. Our study demonstrates that progressing sequencing technology and bioinformatic improvements will bring more knowledge into virome composition, diversity and potentially their important functions.

**Highlights:**

1. Virus-like particles were enriched from human stool samples;
2. Viral nucleotides were sequenced with Oxford Nanopore Technology with and without amplification;
3. Gut virome in humans showed highly individualized composition;
4. Novel sequences and contigs were found to be the majority in the resulted sequences;
5. Epigenetic modifications were detected directly on virus genomes.

## Introduction

The human gut is home to tremendous amount of microbes [1]. They inhabit different ecological niches in the gut, forming complex interaction networks between themselves and with the human cells, and the dynamic balance between gut microbiome and host is required for human health [2–6]. Studies in human cohort and mouse models, among others, have confirmed that gut microbial communities are associated with increasing number of some of metabolism diseases and infectious diseases, providing insights as well as potential targets for future monitoring and therapies [3, 7–14].

The gut microbiome contains bacteria, archaea, fungi, protozoa, and, lastly but not leastly virus. The most abundant cellular members of the microbiome are bacteria and archaea (account for more than 99% of biomass), and have received most attention in human microbiome studies over the years [15–17]. Yet, the advances in next-generation sequencing (NGS) technology and bioinformatic tools have also facilitated the development of human virome studies. Metagenomic analysis suggests that the gut of healthy humans harbors commensal virus, including phages, DNA virus and RNA virus [18–22]. Virome (phages and other host viruses) play roles in intestinal physiology, enteric immune system, host health and disease [23, 24]. The dynamic balance between the virome and the intestinal immune system is finely regulated by cytokines secreted by immune cells [20, 25]. For instance, virome changes in inflammatory bowel disease (IBD) (Crohn’s disease and ulcerative colitis) are disease specific [26]. Phages residing in mucosal surfaces can influence the host by providing nonhost-derived immunity against bacterial infections [26, 27]. By inducing interferons (IFNs), commensal virus can protect from gut inflammation during tissue damage [28, 29]. However, using current short read sequencing technologies, such as Illumina, can only offer knowledge on gut virome that is both biased and fragmentary.

Oxford Nanopore Technologies (ONT) as one of the emerging single-molecule real-time sequencing technology (SMRT) has the advantage of rapid library preparation, ultra-long reads and real-time data acquisition [30–32]. For virome, ONT sequencing has the potential to acquire virus genome by producing genome-length reads that cover all of mutation within a single virus particle. In addition, biological nanopores are able to discriminate not only the genome but also single base modifications such as 5-methylation of cytosine (5mC for DNA and m5C for RNA) and 6-methylation of adenine (6mA for DNA and m6A for RNA) in native DNA/RNA [32]. Increasing evidence in last years suggests that DNA/RNA methylation can influence biological function, including regulation of DNA/RNA replication and repair, and gene expression [33–36]. Recently, *Oliveira* et al. reported that DNA methyltransterase in *Clostridioides difficile* has able to mediate sporulation, *C. difficile* disease transmission and pathogenesis [37]. *Xue* et al. reported viral N^6^-methyladenosine could upregulate replication and pathogenesis of human respiratory syncytial virus [38]. These findings suggest that epigenetic regulation is also important for the pathogenesis of important pathogens.

To profile the gut virome in healthy adults, including identity as well as potential epigenetic information in the virome, we developed a protocol combining physical enrichment, optional reverse transcription and amplification of nucleotides, and bioinformatic analytical pipelines, and firstly characterized the virome in five healthy humans using the ONT PromethION platform. We were able to generate long reads for virome up to tens of kilobases, resulting in many novel contigs that do not have matches in the available databases, and also for abundant virus we could detect epigenetic signals. These discoveries are instructive to future investigations into the genomics, epigenomics and potential function of human gut virome.

## Material and Methods

### 1. Enrichment and purification of virus-like particles (VLPs)

Each frozen faecel samples (approximately 1.5 g) from five individuals who provided written informed consent were resuspended in 15 ml sterile Phosphate Buffered Saline and homogenized thoroughly. The suspension was centrifuged at 4,500 rpm for 10 min at the 4 ℃ to remove large food residues (Beckman Coulter Allegra^TM^ X-22R). Transfer the supernatant to fresh tubes and centrifuged at 4,500 rpm for 10 min at the 4 ℃ again. The suspernatant was filtered through 0.45 μm PVDF membrane (Millipore) to remove eukaryotic and bacterial cell-sized particles before ultracentrifugating at 180,000× g for 3 hours at the 4 ℃ (Beckman Coulter XP-100). The pellets were resuspended in 400 µl sterile Phosphate Buffered Saline and treated with 8 U of TURBO DNaseⅠ (Ambion) and 20 U of RNase A (Fermentas) at 37 ℃ for 30 min. The viral nucleic acids (DNA and RNA) were extracted by using QIAamp MinElute Virus Spin Kit (Qiagen) following the manufacturer’s instructions and eluted into RNase-free water. [39, 40]

### 2. Reverse transcription and random amplification

Viral first strand cDNA was synthesized in a 20 μl reaction mixture with 13 μl of purified vial nucleic acids from each sample and 100 pmol of primer Rrm (5’-GACCATCTAGCGACCTCCAC - NNNNNN-3’), as previously described [39–41]. For the double-strand cDNA synthesis, 100 pmol of primer Rrm and Klenow fragment (3.5 U/μl; Takara) were added. Random amplification was conducted with 8μl of the double-strand cDNA template in a final reaction volume of 200μl, which contained 4μM primer Rm (5’-GCCGGAGCTCTGCAGAATTC-3’), 90 µM dNTPs each, 80 µM Mg^2+^, 10x Buffer and 1 U of KOD-Plus DNA polymerase (Toyobo). The amplification product was purified by agarose gel electrophoresis and QIAquick Gel Extraction Kit (Qiagen).

### 3. PromethION library preparation and sequencing

PromethION library preparation was performed according to the manufacturer’s instructions for the barcoding cDNA/DNA and native DNA (SQK-LSK109 and EXP-NBD104). When multiplexing, all the samples were pooled together. ONT MinKNOW software (v.19.10.1) was used to collect raw sequencing data, and Guppy (v.3.2.4) was used for local base-calling of the raw data after sequencing runs were completed. The PromethION was run for up to 96 h.

### 4. ONT sequence analysis and assembly

Qcat (Oxford Nanopore Technology), python command-line tool for demultiplexing ONT reads from FASTQ files, was used to trim adaptor and barcode sequences. With genomeSize = 2k and default parameters, trimmed raw reads were analysis using Canu v1.9 [42] for virome genome de novo assembly, which includes read correction, read trimming and contig construction.

### 5. Matching to current database

Raw reads qcat trimmed were analysis using mimimap2 [43] to identify gut virome composition, which aligned reads to the reference genome in the National Center for Biotechnology Information (NCBI) virus genome database, including all known viruse genome sequences. To improve the accuracy of the viral taxonomy, two following criteria were adapted: (1) the depth of coverage of reference viral genome >= 5X; (2) the breadth of coverage of the reference viral genome >= 50%.

To assign the taxonomy of assembled contigs by Canu, two approaches and three databases were applied, including minimap2, blastn, NCBI virus genome database, The human gut virome database (GVD) and NCBI nucleotide database. The filter criteria of alignment results of contigs by minimap2 was the same with raw reads, while contig matched length >= 1000 bp with nucleic similarity >= 98% and e-value <= 10^-5^ was adapted as the identified criteria of viruses by blastn.

### 6. Identification and annotation of bacteriophage ORFs

Seeker [44], a new prediction tool via deep learning framework, was used to identify putative phages from contigs in amplified cDNA/DNA group with default parameters. According to the multiPhATE pipeline [45], ORFs in phages were identified by PHANOTATE [46], a tool to annotate phage genomes. Consequently, amino sequences of ORFs were aligned to Phantome (http://www.phantome.org) and pVOGs [47] databases by blastp with parameters “percent of identity >= 60, e-value <= 0.01” and hmm searched to pVOGs by jackhmmer [48] with default parameters, respectly. ORFs repeated in two or more samples were clusterd by usearch [49] with percent of indentity >= 99. To validate these ORFs, we mapped our in-house NGS data using bowtie2 [50] with default parameters. Coverage of ORFs was calculated by weeSAM (https://github.com/centre-for-virus-research/weeSAM) and only ORFs with mapped reads >= 10 were counted.

### 7. Methylation analysis

Tombo v1.5 was used to detect the methylation states of nucleic acids from raw DNA samples [51]. A log-likelihood threshold of 2.5 was used to call methylation and the filter cutoff of methylation sites was defined by the estimated fraction of significantly modified reads >= 0.7 and coverage depth >= 10X. Different methylation sites of 5-methylcytosine (5mC) and N6-Methyladenine DNA Modification (6mA) were visualized by Integrative Genomics Viewer (IGV Version 2.5.3) [52] with default parameters and the putative methyltransferase recognition motifs were identified by MEME (Version 5.0.5) [53] with the following parameters: “-dna –mod zoops”, and webLogo (https://weblogo.berkeley.edu/logo.cgi) was used to plot the logo of the motifs we identified.

### 8. Accession number

The sequencing data were deposited at GSA (Genome Sequence Archive) under BioProject accession no. PRJCA002499. The full protocol is available at https://github.com/caojiabao/VirPipeline.

## Results

### 1. Virome separation, enrichment and sequencing

Since metagenomic sequencing using fecal DNA usually results in only minor fraction of virome sequences, and most of reads will be either from bacteria or archaea, enrichment of viruses are necessary for studying virome in human gut samples. Thus, we have combined a series of enrichment methods including filtration and super centrifugation, to enrich for virus-like-particles (VLPs) in the fecal samples (Figure 1). After VLPs were isolated, additional DNase/RNase treatment were used to remove any potential free-DNA that were not virus-originated. The left-over DNA/RNA were quantified, and half of the nucleotides were directly subjected to ONT DNA library preparation, to profile the DNA virus abundances as well as methylation; and the other half were first subjected to RNA reverse transcription and then amplificaiton with short random primers, to have RNA virus sequenced as cDNA, and improve the chance of low-abundance DNA viruses to be detected in the sequencing results.

**Figure 1.**
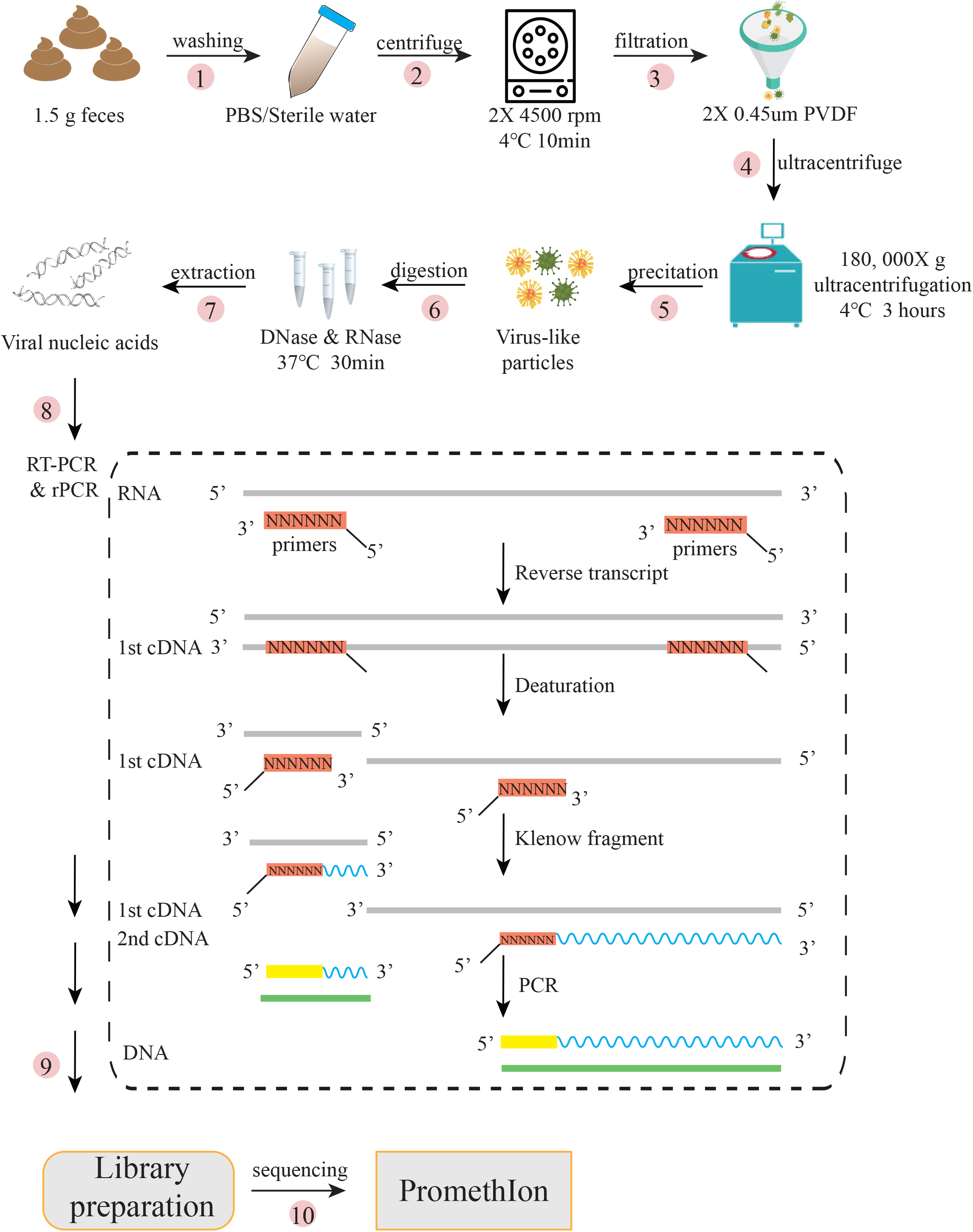
An integrated novel workflow for enrichment of virus-like particles (VLPs), extraction of nucleic acids and ONT sequencing. The complete workflow consists of four fragments: (1) Washing and filtration of faecal samples using PBS/Sterile and PVDF membrane including step 1-3; (2) Precipitation of VLPs including step 4-5; (3) Extraction, amplification and purification of viral nucleic acids including step 6-8; (4) Construction of library and ONT sequencing including step 9-10.

In our study as a primary investigation, we have first profiled five healthy volunteers’ fecal samples with our protocol. Virus-like particle (VLP) fractions of five individuals were enriched, and raw DNA, as well as enriched cDNA/DNA were sequenced using ONT PromthION platform. With one flowcell, the ONT PromthION yielded a total of 8.2 Gb raw data, with a median of 1.7 Gb per sample in amplified group; and 452 Mb raw data, with a median of 67 Mb per sample in raw DNA group (Table S1).

### 2. Virome composition in healthy individuals revealed by ONT sequencing

With sequencing reads from amplified cDNA/DNA results, we have mapped the ONT reads onto NCBI virus genomes and then analyzed the composition across five individuals. Consistent with other studies, our result from amplified cDNA/DNA of virome showed that bacteriophage families were the most frequently detected and accounted for the majority of the intestinal virome in number. The final catalogue of bacteriophages included the *Caudovirals* order (families *siphovirdae*, *podoviridae*), family *Inoviridae* and family *Microviridae.* Meanwhile, the eukaryotic CRESS-DNA viruses (family *genomoviridae*) was also detected (Figure 2; Table S2). Of special note was the presence of the RNA plant viruses including family *Virgaviridae* and *Alphaflexiviridae*, which showed good agreement with the findings of Shkoporov *et al* [54]. We observed that individual specificity is probably a feature of the faecal viral communities (Figure 2), which had been already demonstrated by several previous studies [55–57]. However, uncultured phage WW-nAnB strain 3 belonging to family *inoviridae* was detected to be presented in amplified cDNA/DNA among five individuals.

**Figure 2.**
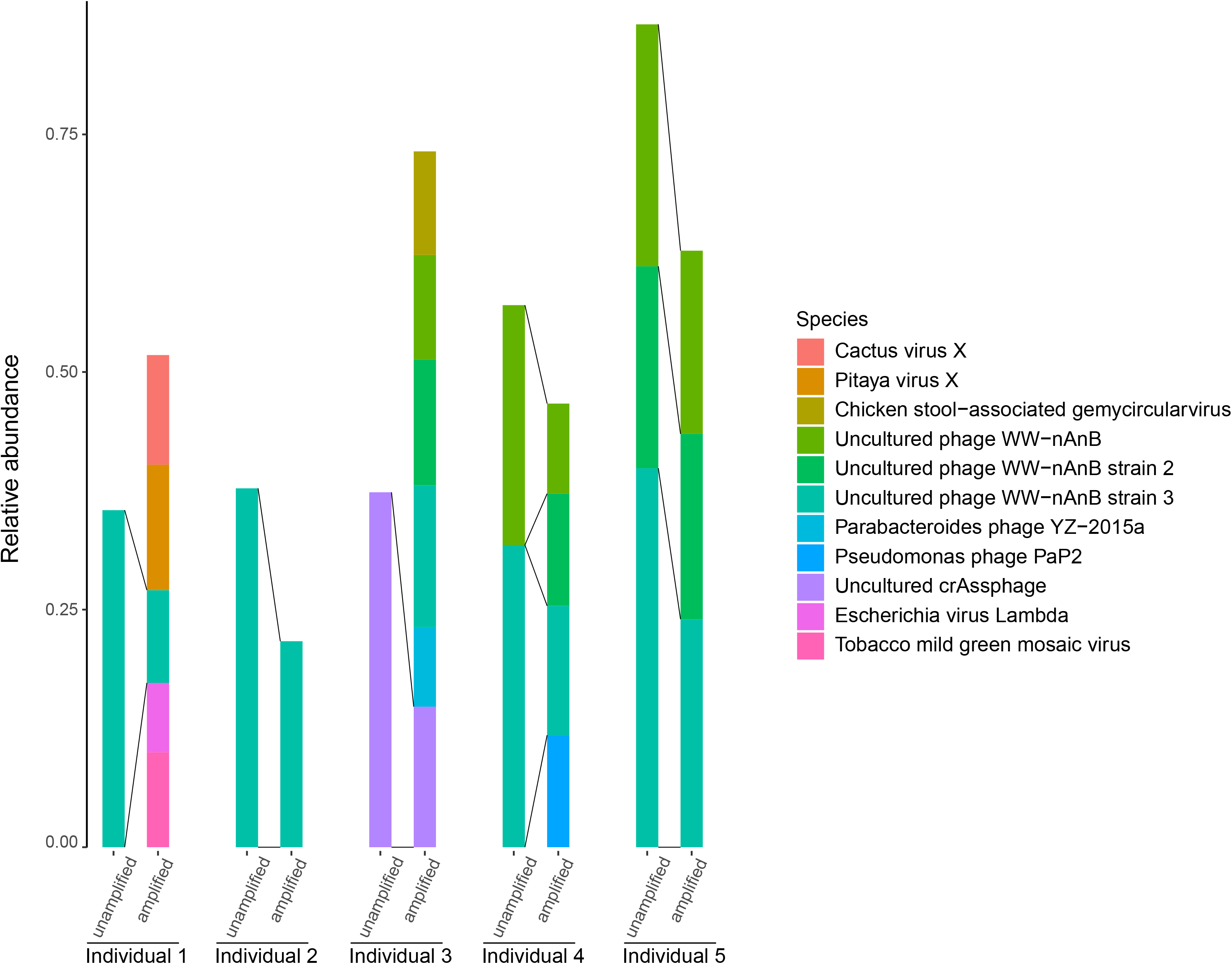
Composition and relative abundance of viruses in each individual. Different color of the bar represents different viral species or strains. The line between unamplified and amplified group in each individual represents common virus species between two groups.

To characterize the potential biases regarding virome composition caused by amplification, we have compared the results from amplified cDNA/DNA and raw virome DNA. We could not compare raw RNA due to the fact that it is still not yet possible to multiplex RNA samples on ONT platforms. The relative abundance of virus each virome was defined using relative proportion of each virus in terms of breadth of coverage on the assembled genome, similar to the definition of bacterial abundances in metagenomic studies. As expected, numbers of viruses are higher in amplified cDNA/DNA results, except for individual 2 and individual 5 who remained the same in terms of virus diversity. Further, abundances of the common existing viruses between raw DNA and amplified cDNA/DNA showed essential shifts in all five individuals, demonstrated detectable bias of reverse transcription as well as random amplification approach we adapted.

### 3. Virome assembly using ONT sequences

We next focused on the assembly of nanopore sequencing reads in five individuals. Assembler Canu was used to assemble the virome sequences separately from raw DNA and amplified cDNA/DNA groups into contigs, which yielded a total of 1564 contigs, with a median of 15 and 347 contigs per sample for raw DNA and amplified cDNA/DNA group. Average length of contigs from raw DNA group was longer than those from amplified cDNA/DNA group (Figure S1). The contigs vary largely in length, ranging from 1kb to 53kb (Figure S2). Consequently, we obtained the identity of certain contigs by matching with NCBI virus genomes, human gut virus database (GVD) [58] and NCBI nucleotide databases, however all with very low matching rate, suggesting a large collection of potentially novel genomes in our results (Table S3). Thus, Seeker was used to identify bacteriophages in amplified cDNA/DNA group. As a result, more than 50% of contigs per sample were identified as phages in amplified cDNA/DNA group. To characterize these bacteriophages, we performed phages ORFs prediction and functional annotation, which yielded average 7 ORFs per contig, ranging from 6 to 10, and average 3 ORFs per kb in length. In addition, ORFs repeated in two or more samples were validated by mapping to additional Illumina Hiseq sequencing data of metagenomics from same samples. We founded that over half ORFs (59.3%, 35/59) can be matched, indicating that these phages stably exist in our data (Table S4).

To examine the potential of covering full virus genome of long sequence produced from the oxford nanopore technology sequencing, we mapped raw reads to the contigs by canu separated from amplified cDNA/DNA and raw DNA. We aggregated and counted the length of raw reads aligned to the longest contigs of our choice from these two groups, and calculated the proportion of raw reads to contigs in length (Figure 3). In amplified VLP cDNA/DNA group, the max value of reads length in each sample was all more than 15% of contigs in length, the highest achieving 40% of contigs’ full length in individual 4 (Figure S3). In raw DNA group, this proportion was higher in general, understandably due to the fact random amplification usually can not reach full length available in the raw DNA. A small number of reads were longer than the final contigs (Figure 3), for which the possible reason for this result was that some long reads were trimmed by canu during assembly, due to the lower quality of part of the sequences being abandoned by Canu.

**Figure 3.**
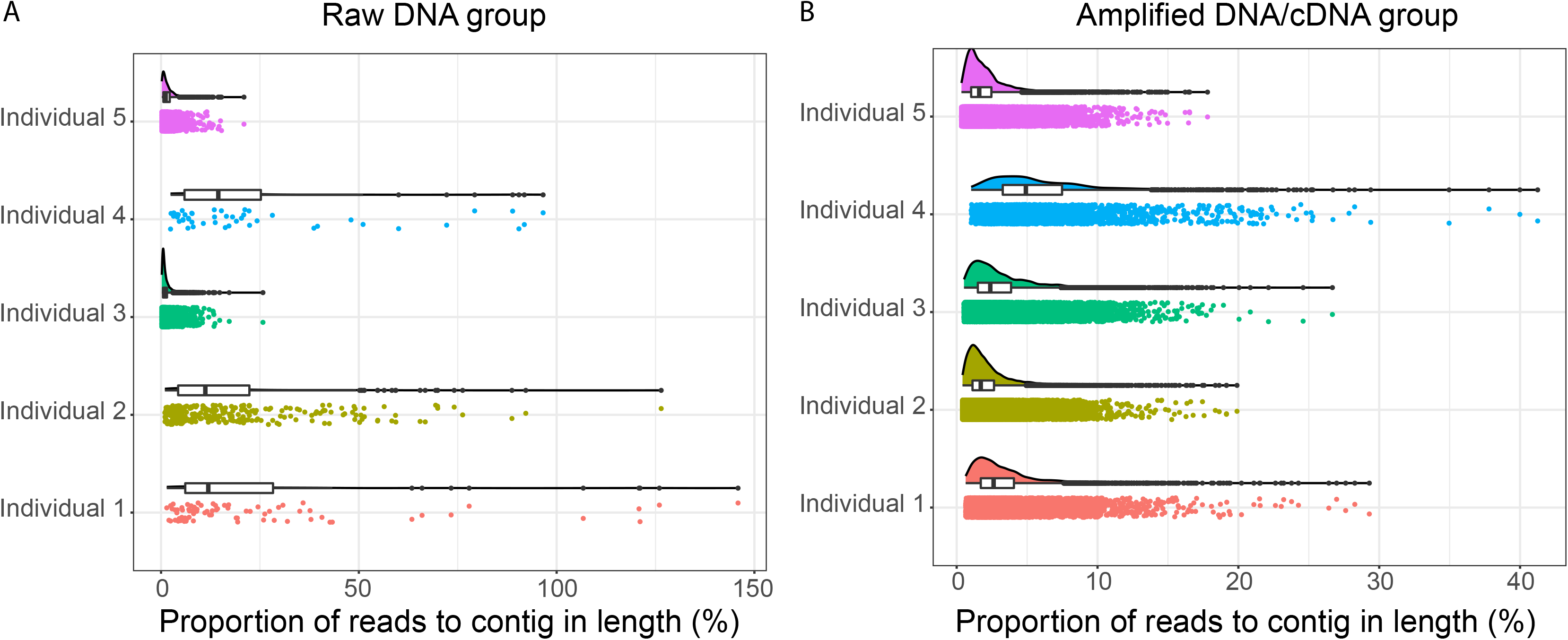
Proportion of raw reads to contigs in length. The proportion of raw reads to contigs in length is shown by combination of scatter, diagram, boxplot and violin chart. Different color represents five different individuals. (A) raw DNA group, (B) amplified DNA/cDNA group.

### 4. Viral epigenomic detection using ONT

In addition, methylation states of bases in DNA from reads can be detected directly by the Oxford Nanopore sequencing without extra laboratory techniques. In this study, we have analyzed methylation signals on a few contigs that reached >10X coverage in raw DNA data in any of individuals. Methylation detection can not be carried out on cDNA and amplified DNA samples, for they will lose all the methylation signals. For the only one of the contigs (contig00000015) with known identity (Uncultured crAssphage), we detected in total 17 5mC and 120 6mA methylation sites covered in the 8kb genome (Figure 4). For 5mC and 6mA methylation, the nucleotide motifs YCHYTTA****C****TWMRECT (e-value = 1.2 x 10^-2^) and motif MADWDTW****A****NADYYWW (e-value = 2.5 x 10^-4^) were identified respectively, with the methylated nucleotide highlighted in bold italic.

**Figure 4.**
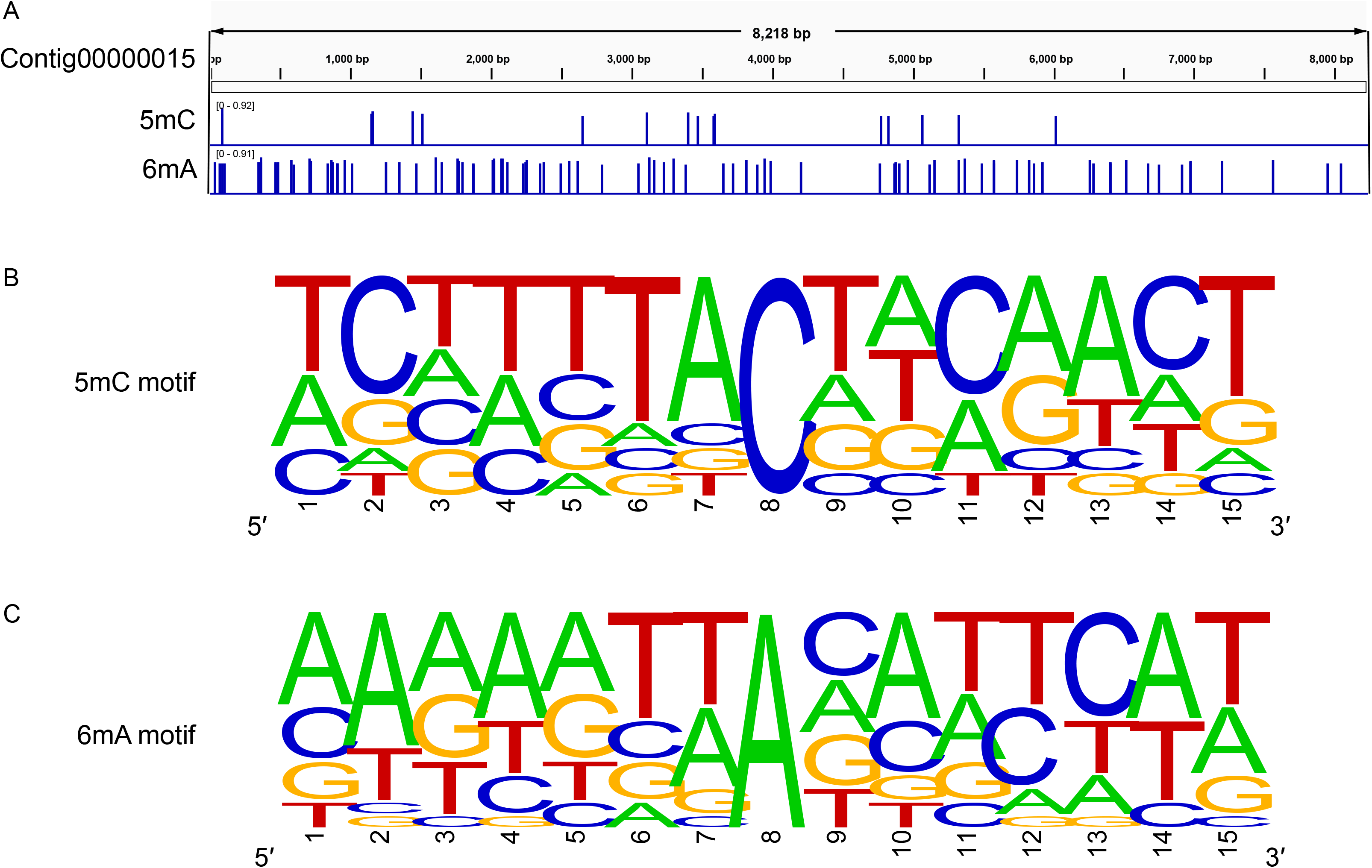
Different methylation sites identification and motif recognition of viral contig. (A) the distribution of methylation sites (5mC and 6mA) in the contig00000015; (B) 5mC motif; (C) 6mA motif.

We have also found another 4 contigs with >10X coverage, but they do not have detectable relative in the databases we have searched against. They are potentially novel viruses and very unlikely to be bacteria or archaea, as our protocol has removed most of the none VLPs, and bacteria/archaea contigs would most likely have matches in the databases we have searched. They have 5mC methylation sites ranging from 0.3 to 1.8 (% of genome) and 6mA methylation sites ranging from 0.7 to 2.5 (% of genome). The motifs for methylation are also extremely diverse, including NHHYYKG***C***DHNNN, WDWADDW***C***DWYNDDW, MNNNNTR***C***GBNNNND and HDNBYDV***C***VVVVNNH for 5mC, and WHWHNYD***A***HNNYYHTT, VNWWDWH***A***YBYNNNT, DRNVRKK***A***BBNDNNN and NRNARND***A***SYAHHNH for 6mA (Figure S4). Since most of the methylation studies are performed in eukaryotes, and only starting in bacteria with limited information available, it is yet difficult to compare the methylation profiles to understand its underlying mechanisms.

## Discussion

Our study combined the physical enrichment of VLPs in fecal samples, nucleotide amplification with the latest sequencing technology to establish a complete workflow of human gut virome profiling. With longer reads as well as richer information of additional epigenetic modifications, developments in sequencing technology could bring another round of revolution in metagenome as well as virome investigations. More importantly, despite the complex steps before sequencing were designed for maximum enrichment of VLPs, as well as removal of any DNA/RNA that were not of virus origin, and thus inevitably makes it relatively time-consuming, ONT could carry out sequencing and produce reads nearly simultaneously, making it possible to finish data generation from samples with five days of working time, and potentially even shorter if PromethION was run for less time, or data were analyzed during the course of being generated (real-time); comparatively, Illumina based platforms usually takes longer to generate enough read length for downstream analyses. There are several studies who have already utilized this property for fast pathogen detection in infectious diseases [59–61], there are cases of virome analysis that might require such time efficiency as well, and our protocol provides a feasible choice for virome studies that also requires short turn-around time.

We also compared the effect of reverse transcription and consequent amplifications on virome analysis with ONT. Such steps were used for several reasons.

Firstly, despite ONT is capable of sequencing DNA or RNA directly, multiplexing is still only possible for DNA libraries, and directly sequencing RNA is not yet cost-effective for virome analysis, while cDNA is a better alternative. Also, to utilize the capacity of sequencing on ONT, very high concentrations of DNA or RNA libraries are required to generate enough reads, yet this is also very difficult for virome DNA/RNA, who usually only reaches 10% of the required input from 1.5 grams of fecal samples. Amplification does lead to higher amount of reads and enables detection of low abundance viruses to be detected, as revealed in our study; but it also leads to on average shorter reads and certain biases in the estimations of virus abundances, resulting from both affinity to random primers as well as PCR produced artefacts. Lastly, amplified cDNA/DNA loses all the methylation modifications on the viral nucleotides and prevents investigations into this potentially vital epigenetic information. Thus, future investigator needed to balance the pros and cons of amplification processes, and could take advantage of our approach of using both raw DNA (and/or RNA) and amplified cDNA/DNA for sequencing, gaining complementary information within the same sequencing run.

The profiles of virome in our studied individuals suggest a highly diverse virome, with only small amount of “core” viruses shared in between. This core could shrink even further with increasing number of individuals while the total diversity of viruses increase, which we plan to investigate in the future. We found phages making the majority part of the virome in the human gut, plus a few host viruses; while the mystery of plant RNA viruses is again present with ONT data, many reads achieving > 1kb in length, whether they are left-overs from our plant-based food, or rather human viruses with their closest relatives in the plant-associated viruses, still call for more investigations, especially within functional experiments. Our data suggest that there are potentially high number of unknown, novel viruses in the human gut, as our assembled contigs have very low rate of matches in the current databases; we consider those not likely to be contaminations due to our vigorous depletion of any non-VLPs and non-viral nucleotides, plus the fact that they do not have any match in nucleotide collection of NCBI either.

It’s also the first time we demonstrate that the phage genomes are methylated via direct sequencing. RSV viruses and influenza viruses are known to have m6A methylation on their RNA genome [38, 62], detected with more complex methods with low throughput, while DNA phages (or other DNA viruses) are not yet studied to our knowledge. In *E. coli* and *C. difficile* it is shown that 6mA is the main form of DNA methylation, while eukaryotes usually lack this form, and in our results phages also have 6mA as the main form of DNA methylation [37, 63, 64]. Since DNA methylations play an important role in bacterial defense against phages, how phage genome becomes methylated, and the consequent impact on phage life cycle and interactions with bacterial hosts remain to be explored with dedicated studies. Besides, it remains possible the motifs of methylation between phage and bacteria are intrinsically linked, and provide additional information to determine the host range of phages; this would require increasing the current knowledge on epigenetics of both bacteria and phages in the future.

## Conclusions

To summarize, we developed and pilot-tested a thorough protocol for human gut virome analysis using the lastest ONT sequencing platform, and generated novel insights into the individuality, diversity of gut virome with new sequencing data. Our approach of course can be applied for other virome studies, including animal gut, soil and water virome etc., and accumulating both sequences as well as epigenetic information on those samples, have the long potential of opening up new directions in metagenomic, microbiological and medical researches.

## Acknowledgements

This work was supported by the National Key Research and Development Program of China (grant number 2018YFC2000500), the Strategic Priority Research Program of the Chinese Academy of Sciences (grant number XDB29020000), and the National Natural Science Foundation of China (grant number 31771481 and 91857101).

## Notes

### Competing Interest Statement

The authors have declared no competing interest.

