## Supplementary Figures for "Profiling of Human Gut Virome with Oxford Nanopore Technology"

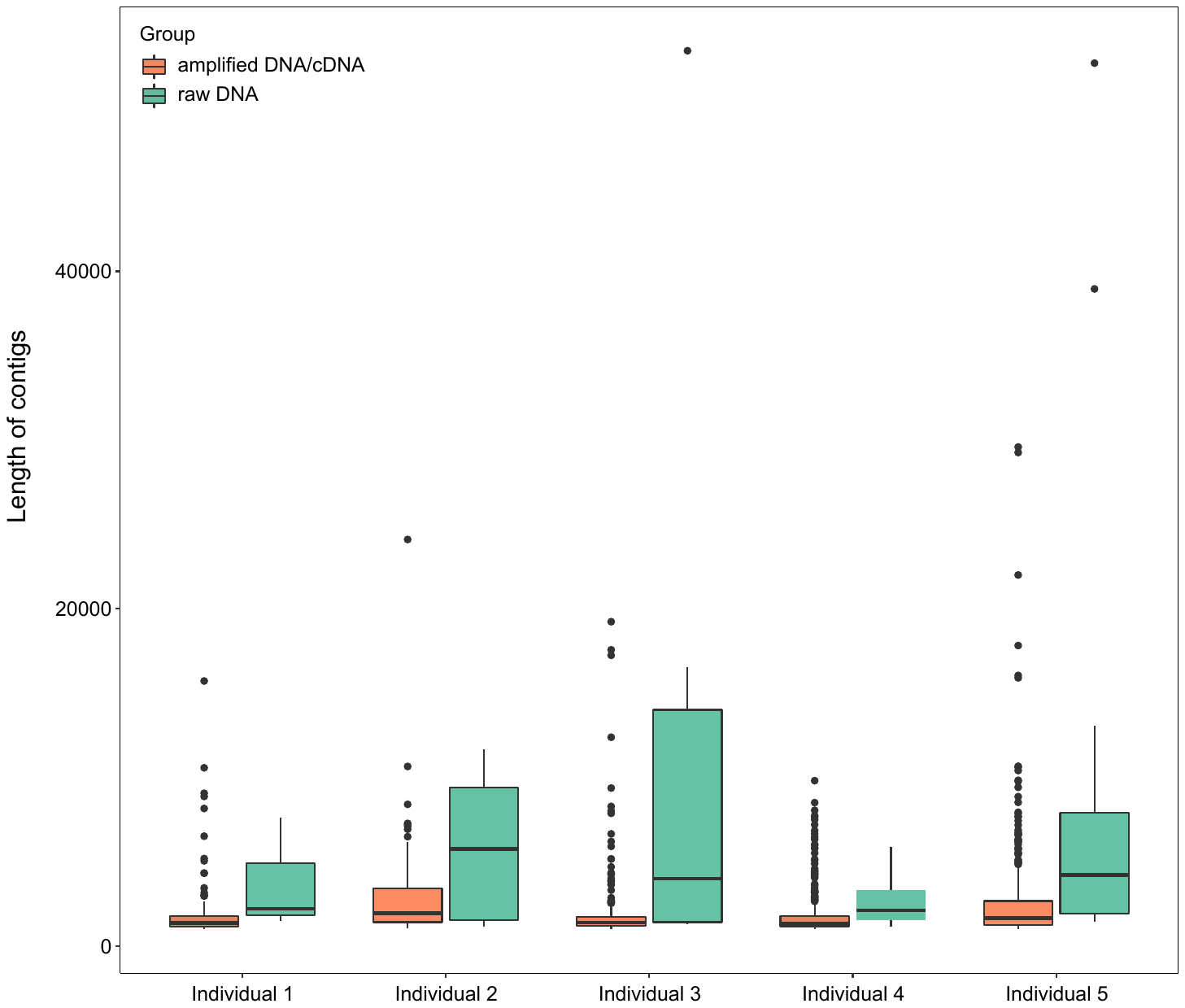

**Supplementary figure 1. Comparison of contig length from amplified DNA/cDNA and raw DNA samples.**

The comparison of contigs length from amplified DNA/cDNA and raw DNA samples is shown. Orange box represents contigs from amplified DNA/cDNA samples and blue box represents contigs from raw DNA samples.

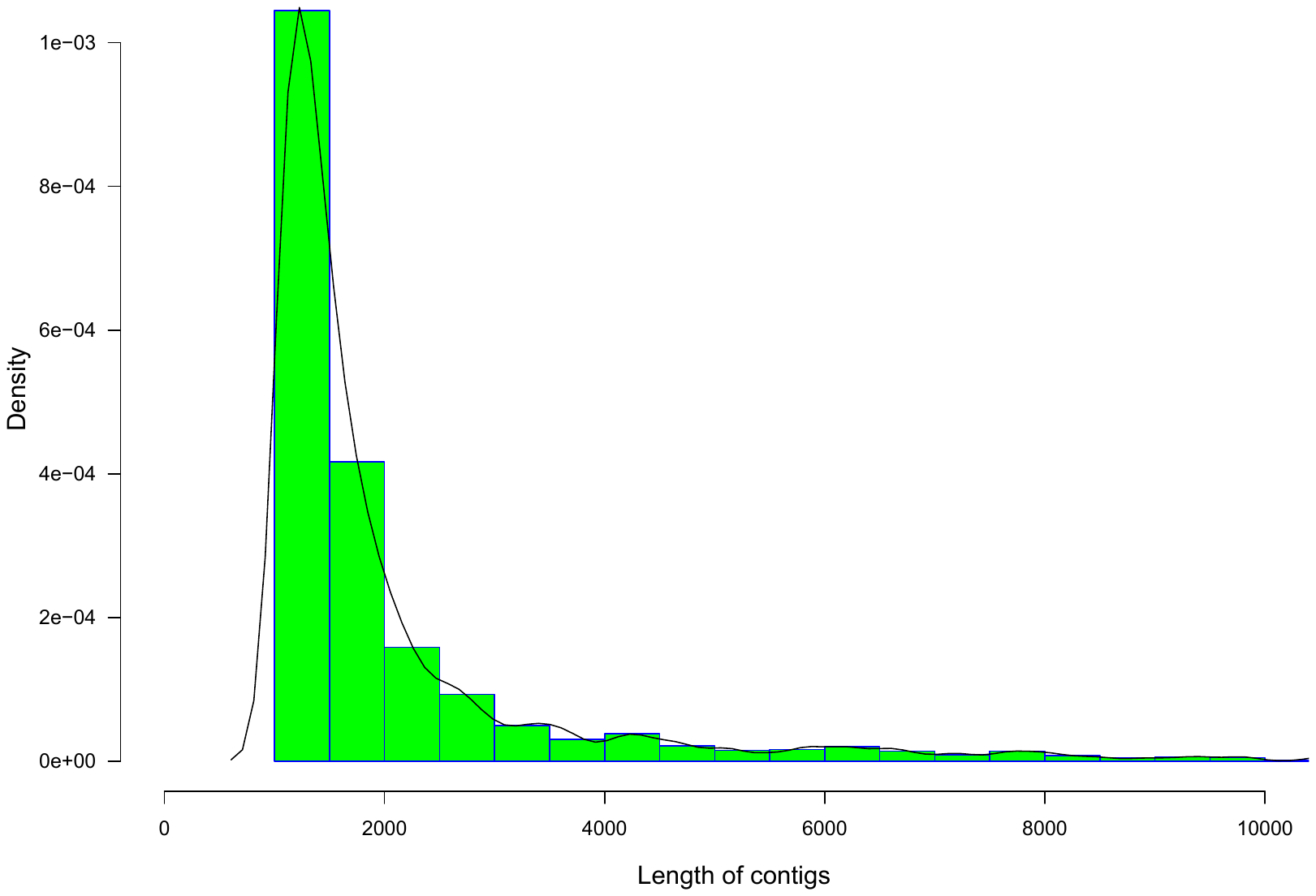

**Supplementary figure 2. Histogram and density of contig length distribution from five individuals.**

Histogram and density of total 1,564 contigs’ length from five individuals are shown. The contigs vary largely in length, ranging from 1kb to 53kb, with a median of 1,469bp and mean of 2,289bp.

**Supplementary figure 3. Comparison of qualified reads length from amplified DNA/cDNA and raw DNA samples.**

Qualified read length of amplified DNA/cDNA and raw DNA samples are shown in boxplot. Orange represents amplified DNA/cDNA sample and green represents raw DNA sample. Qualified read length ranges from 100bp to 24,039bp in amplified DNA/cDNA, from 100bp to 49,728bp in raw DNA.

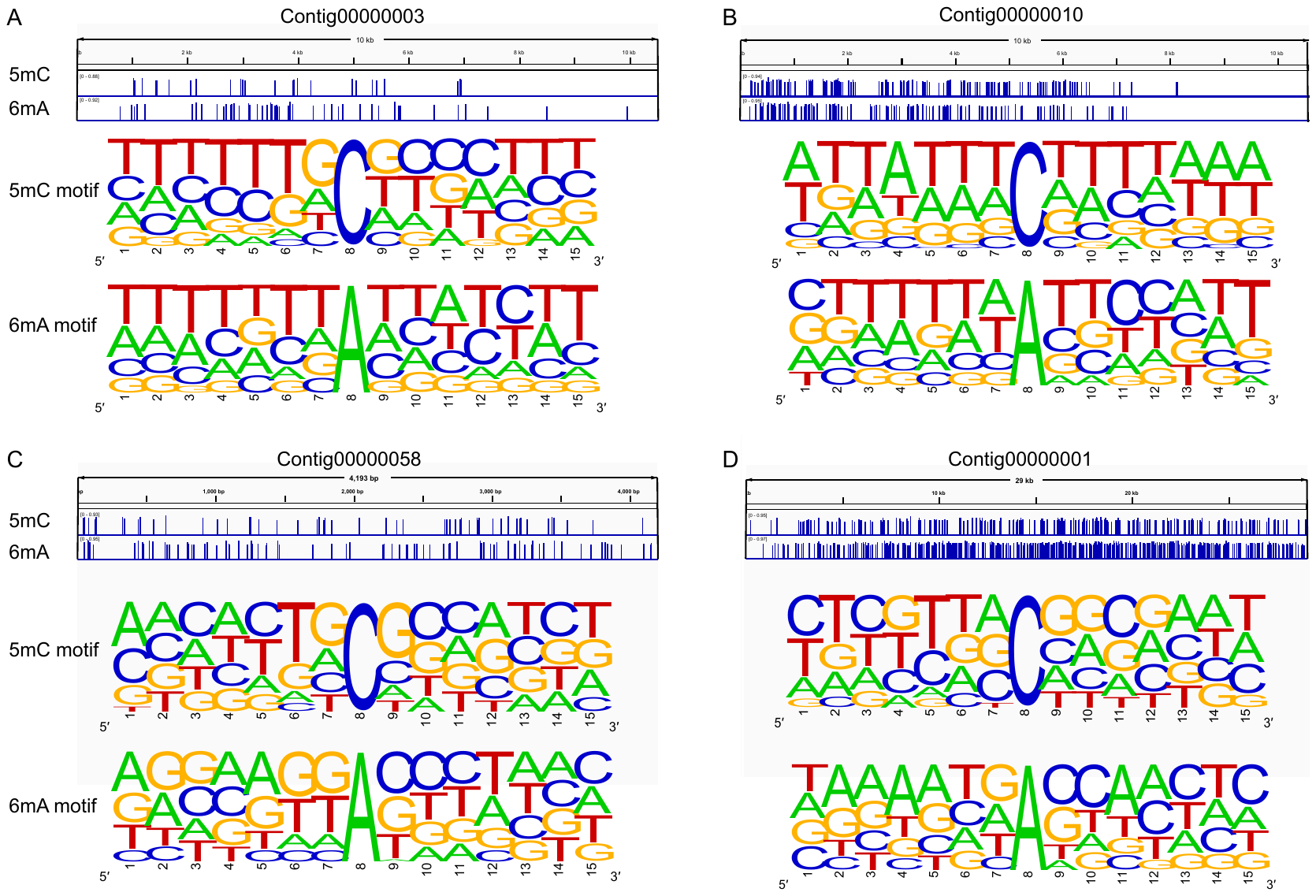

**Supplementary figure 4. Methylation sites identification and motif recognition of four viral contigs.**

5mC and 6mA methylation sites distribution in Contig00000003 (A), Contig00000010 (B), Contig00000003 (C) and Contig00000001 (D) are shown in IGV and their motifs of different methylation model are recognized.
