## Supplementary Tables for "Profiling of Human Gut Virome with Oxford Nanopore Technology"

**Supplementary table 1**: Extraction of nucleic acids and output of PromethION sequencing

| Individual | Group | DNA/cDNA (ng) | DNA/cDNA (ng) after TR | Input (ng) | Output (Mb) |
| --- | --- | --- | --- | --- | --- |
| 1 | raw DNA | NA | 100.8 | 1185.6 | 77 |
| 2 | raw DNA | NA | 107.52 |  | 48 |
| 3 | raw DNA | NA | 50.4 |  | 57 |
| 4 | raw DNA | NA | 263.04 |  | 203 |
| 5 | raw DNA | NA | 276.48 |  | 67 |
| 1 | amplified DNA/cDNA | 2,100 | 5,664 |  | 1,700 |
| 2 | amplified DNA/cDNA | 2,570 | 8,736 |  | 892 |
| 3 | amplified DNA/cDNA | 1,700 | 5,808 |  | 1,500 |
| 4 | amplified DNA/cDNA | 2,130 | 6,672 |  | 2,200 |
| 5 | amplified DNA/cDNA | 1,200 | 6,960 |  | 1,900 |

Note: the column 3 represents total nucleic acids before library construction. The column 4 represents total nucleic acids after terminal repair (TR). When multiplexing, all the barcoded samples were pooled together.

**Supplementary table 2:** The identification of viral family and species in gut

|  |  | Individual 1 | Individual 2 | Individual 3 | Individual 4 | Individual 5 |
| --- | --- | --- | --- | --- | --- | --- |
| Family | Inoviridae | yes | yes | yes | yes | yes |
|  | Siphoviridae | yes |  |  |  |  |
|  | Genomoviridae |  |  | yes |  |  |
|  | Podoviridae | yes |  | yes | yes |  |
|  | Microviridae |  |  | yes |  |  |
|  | Alphaflexiviridae | yes |  |  |  |  |
|  | Virgaviridae | yes |  |  |  |  |
| Species | Cactus virus X | yes |  |  |  |  |
|  | Pitaya virus X | yes |  |  |  |  |
|  | Uncultured phage WW-nAnB strain 3 | yes | yes | yes | yes | yes |
|  | Lambdavirus,Escherichia virus Lambda | yes |  |  |  |  |
|  | Tobacco mild green mosaic virus | yes |  |  |  |  |
|  | Chicken stool-associated gemycirc |  |  | yes |  |  |
|  | uncultured crAssphage |  |  | yes | yes | yes |
|  | uncultured phage WW-nAnB |  |  | yes | yes | yes |
|  | Uncultured phage WW-nAnB strain 2 |  |  | yes |  |  |
|  | Parabacteroides phage YZ-2015a |  |  | yes |  |  |
|  | Pseudomonas phage PaP2 |  |  |  | yes |  |

Note: character 'yes' represents the existence of this virus in the individual

**Supplementary table 3**: The number of identified certain contigs

| Individual | Group | NCBI (minimap2) | GVD (minimap2) | NCBI NT (Blastn) |
| --- | --- | --- | --- | --- |
| 1 | raw DNA | 0 | 0 | 0 |
| 2 | raw DNA | 0 | 0 | 0 |
| 3 | raw DNA | 0 | 1 | 8 |
| 4 | raw DNA | 0 | 0 | 0 |
| 5 | raw DNA | 0 | 0 | 0 |
| 1 | amplified DNA/cDNA | 0 | 3 | 5 |
| 2 | amplified DNA/cDNA | 0 | 1 | 0 |
| 3 | amplified DNA/cDNA | 1 | 2 | 10 |
| 4 | amplified DNA/cDNA | 0 | 2 | 0 |
| 5 | amplified DNA/cDNA | 0 | 5 | 0 |

Note: NCBI, NCBI viruses genome database; GVD, The human Gut Virome database; NCBI NT, NCBI nucleotide database

**Supplementary table 4**: ORFs information mapped by Illumina data

| ORF | Length (nt) | Occurrence in samples | Average coverage (%) | Average depth | Annotation |
| --- | --- | --- | --- | --- | --- |
| orf1 | 273 | Individual_5 | 100.0 | 449.6 | hypothetical protein SWRG_00180 [Synechococcus phage S-RIM2 R21_2007] |
| orf2 | 1296 | Individual_2 Individual_5 | 82.0 | 379.7 | NA |
| orf3 | 150 | Individual_2 Individual_5 | 100.0 | 263.0 | NA |
| orf4 | 198 | Individual_2 Individual_4 Individual_5 | 100.0 | 213.2 | hypothetical protein [Bacillus phage PM1] |
| orf5 | 171 | Individual_2 Individual_4 Individual_5 | 100.0 | 176.3 | putative dna polymerase [Bacillus phage CP-51] |
| orf6 | 138 | Individual_2 Individual_4 Individual_5 | 100.0 | 80.0 | NA |
| orf7 | 222 | Individual_2 Individual_4 Individual_5 | 100.0 | 347.3 | putative superinfection exclusion protein [Serratia phage phiMAM1] |
| orf8 | 189 | Individual_2 Individual_4 Individual_5 | 100.0 | 196.7 | putative scaffold protein [Streptococcus phage T12] |
| orf9 | 294 | Individual_2 Individual_4 Individual_5 | 100.0 | 371.5 | putative phage tail fiber protein [Celeribacter phage P12053L] |
| orf10 | 1659 | Individual_1 Individual_2 Individual_3 Individual_4 Individual_5 | 91.0 | 6595.6 | NA |
| orf11 | 264 | Individual_1 | 100.0 | 72.9 | aerobic ribonucleoside-diphosphate reductase small subunit [Escherichia phage APCEc03] |
| orf12 | 111 | Individual_5 | 99.1 | 6.5 | NA |
| orf13 | 141 | Individual_1 | 100.0 | 13.4 | NA |
| orf14 | 222 | Individual_4 | 100.0 | 16.3 | hypothetical protein SWZG_00267 [Synechococcus phage S-SKS1] |
| orf15 | 342 | Individual_4 | 99.4 | 21.4 | putative major capsid protein [Gokushovirinae GNX3R] |
| orf16 | 189 | Individual_4 | 99.5 | 9.1 | NA |
| orf17 | 90 | Individual_5 | 100.0 | 34.7 | NA |
| orf18 | 180 | Individual_1 Individual_2 Individual_4 Individual_5 | 100.0 | 4853.4 | hypothetical protein Lu11_0066 [Pseudomonas phage Lu11] |
| orf19 | 240 | Individual_1 Individual_2 Individual_5 | 100.0 | 4596.0 | hypothetical protein phiST2_0181 [Vibrio phage phi-ST2] |
| orf20 | 378 | Individual_2 Individual_4 Individual_5 | 97.2 | 882.2 | NA |
| orf21 | 558 | Individual_5 | 100.0 | 184.5 | hypothetical protein B40-8008 [Bacteroides phage B40-8] |
| orf22 | 450 | Individual_5 | 100.0 | 762.5 | hypothetical protein rtp67 [Escherichia virus Rtp] |
| orf23 | 120 | Individual_5 | 99.2 | 9.1 | [Cellulophaga phage phi10:1]; |
| orf24 | 399 | Individual_5 | 100.0 | 903.1 | hypothetical protein [Delftia phage IME-DE1] |
| orf25 | 198 | Individual_5 | 100.0 | 557.8 | NA |
| orf26 | 144 | Individual_5 | 100.0 | 272.2 | gp131 [Brochothrix phage A9] |
| orf27 | 237 | Individual_5 | 100.0 | 105.4 | endolysin [Bacillus phage phi4I1] |
| orf28 | 396 | Individual_1 Individual_2 Individual_4 Individual_5 | 100.0 | 4932.0 | conserved hypothetical protein [Edwardsiella phage PEi26] |
| orf29 | 348 | Individual_5 | 100.0 | 189.7 | gp67 [Mycobacterium phage Konstantine] |
| orf30 | 384 | Individual_5 | 100.0 | 206.2 | hypothetical protein GBK2_35 [Geobacillus phage GBK2] |
| orf31 | 210 | Individual_4 | 100.0 | 13.5 | NA |
| orf32 | 306 | Individual_5 | 100.0 | 482.8 | NA |
| orf33 | 105 | Individual_5 | 99.1 | 6.9 | NA |
| orf34 | 282 | Individual_5 | 100.0 | 13.4 | NA |
| orf35 | 789 | Individual_4 Individual_5 | 97.2 | 2207.5 | NA |

Note: “Average coverage presents” represents the percent of sites in the genome which have coverage, “Average depth” represents mean read depth. The column of “Annotation” represents the best hit from blast and hmm search.
